# Lower Levels of Task-Induced Hippocampal Activation Predict Better Performance on a Separate Verbal Memory Evaluation

**DOI:** 10.1101/2025.06.02.656665

**Authors:** Caitlin A. Sisk, Kathleen B. McDermott, Jed T. Elison, Meghan A. Swanson, Gagan S. Wig, Adrian W. Gilmore, Steven M. Nelson

## Abstract

Inter-individual differences in learning-related activation in the hippocampus and its subregions predict specific differences in memory abilities. The complexity of these relationships creates a setting that is ripe for theoretically informative investigation, but that can also yield reports of spurious relationships that do not reflect underlying neurobiological associations. Across-study replication is therefore a critical step toward understanding how individual differences in hippocampal activity drive individual differences in memory ability. In the domain of verbal memory, Wig et al. (2008) identified a negative relationship between task-induced activation in right posterior HPC (pHPC) and out-of-scanner verbal memory test performance. Replicating this result in an independent sample of 86 participants, we identified the same negative correlation between pHPC activation during an in-scanner Lithuanian-English paired associate learning task and out-of-scanner California Verbal Learning Test (CVLT-II) performance. In a follow-up exploratory voxel-wise association analysis, we observed the same negative correlation between CVLT-II performance and task-induced activity across three regions of the medial temporal lobe: the left perirhinal/entorhinal cortex, right parahippocampal gyrus, and left posterior hippocampus. This replication and accompanying extension represent a critical step toward understanding how pHPC supports verbal memory by answering basic questions concerning the reliability and selectivity of this previously identified relationship.

## Introduction

The hippocampus, seated in the medial temporal lobe, plays a key role in human and non-human animal memory function [1,2]. Although this basic observation has long been appreciated (e.g., [3]), functional dissociations between anterior and posterior aspects of the structure only became major targets of research sometime later (see [4] for review). Thus, while “the” role of the hippocampus in memory has been the study of psychologists and neuroscientists for many decades, investigations into the nature of how different parts of the hippocampus might differentially contribute to memory or other cognitive processes remain ongoing. A consensus seems to be emerging that anterior aspects of the hippocampus tend to process more abstract information, whereas posterior aspects tend to process fine-grained details (see [5]).

Given its association with fine-grained information, a natural prediction may be that greater activity during task-based periods, compared to periods of rest, might predict a stronger general memory ability. However, evidence suggests the opposite: less activity in the posterior hippocampus during active encoding conditions may instead be a better index of memory performance. Across two studies, Wig et al. [6] identified a negative correlation across subjects between task-induced posterior hippocampal activation and performance on out-of-scanner testing of verbal memory. Critically, neither in-scanner task used in Wig et al. [6] required explicit encoding demands. Rather, they involved odd-even number judgements or responding to onsets and offsets of a flickering checkerboard. The authors suggested that, because their analyses compared task activation to a resting-state baseline, consolidation-related mechanisms captured during rest could likely account for their pattern of results.

The hypothesis forwarded by Wig et al. [6] would suggest that similar negative correlations should be observed between task-states and out-of-scanner memory performance in different task conditions. In this report, we re-analyze data from a large sample of participants (N = 86) in which individual differences in general learning ability were previously reported [7] based on rates of learning Lithuanian-English word pairs. Using these data, we ask if any correlation between task-activation and out-of-scanner learning might be obtained. As the data from Nelson et al. [7] involved an explicit encoding of associations, the task conditions differ substantially from those reported by Wig et al. [6] and therefore constitute a strong test of the general mechanisms hypothesized in that paper. A critical second question concerned the localization of these effects: if any were observed, would they fall specifically in the posterior hippocampus, as in Wig et al. [6], or would they be located across the hippocampus or medial temporal lobe more broadly?

## Methods

### Participants

This study involved a re-analysis of data from participants initially reported in Nelson et al. [7]. One hundred participants were recruited via Craigslist (www.craigslist.org) and community-posted flyers in the St. Louis area. Of the 14 excluded participants, 6 were excluded for excessive movement, 6 for failure to comply with task instructions (failure to keep eyes open or failure to respond), 1 for failure to complete the experiment, and 1 for opting out of participation due to illness. This resulted in a final sample of 86 participants (36 female) between the ages of 18 and 31 years (*mean*: 24.82) who had completed between 10 and 22 years of education (*mean*: 15). Inclusion criteria included right-handedness, English as a native language, and normal or corrected-to-normal vision. Exclusion criteria included reported history of neurological or psychiatric illness. All participants were consented according to the guidelines of Washington University’s Human Research Protection Office and were compensated $25 per hour of participation.

### Stimuli

Stimuli consisted of 45 direct Lithuanian–English translations (e.g., AKIS–EYE) of concrete nouns from published norms [8] presented in all capital letters with no typographic ligatures or diacritical marks. Stimuli were presented in white 48-point Arial typeface on a black background. Lithuanian words varied in character length from 4 to 9 (*mean*: 5.96), and English words varied in character length from 3 to 8 (*mean*: 4.56). Lithuanian words varied in number of syllables from 2 to 4 (*mean*: 2.4), and English words varied in number of syllables from 1 to 2 (*mean*: 1.22). Participants reported no prior knowledge of the Lithuanian language.

### Procedure

The experimental procedure consisted of four phases across two sessions of a Lithuanian-word-learning task. Phases 1-3 took place during Session 1, and phase 4 took place 2 days later during Session 2 (Figure 1). The first phase, Study 1, took place inside of the scanner during active collection of blood oxygen level-dependent (BOLD) response. In Study 1, participants were shown 45 Lithuanian–English word pairs one at a time in random order and were instructed to memorize the word pairs. Participants were told that a later recall test would occur in which they would be shown the Lithuanian cue and would be required to provide the matching English translation. During Study 1, each Lithuanian–English word pair was presented for 3.5s with jittered interstimulus intervals of 1.5-6.5s. BOLD response was again collected during the Study 2 portion of the experiment. Results reported here utilize fMRI data collected during the Study 1 portion of the experiment.

**Figure 1.**
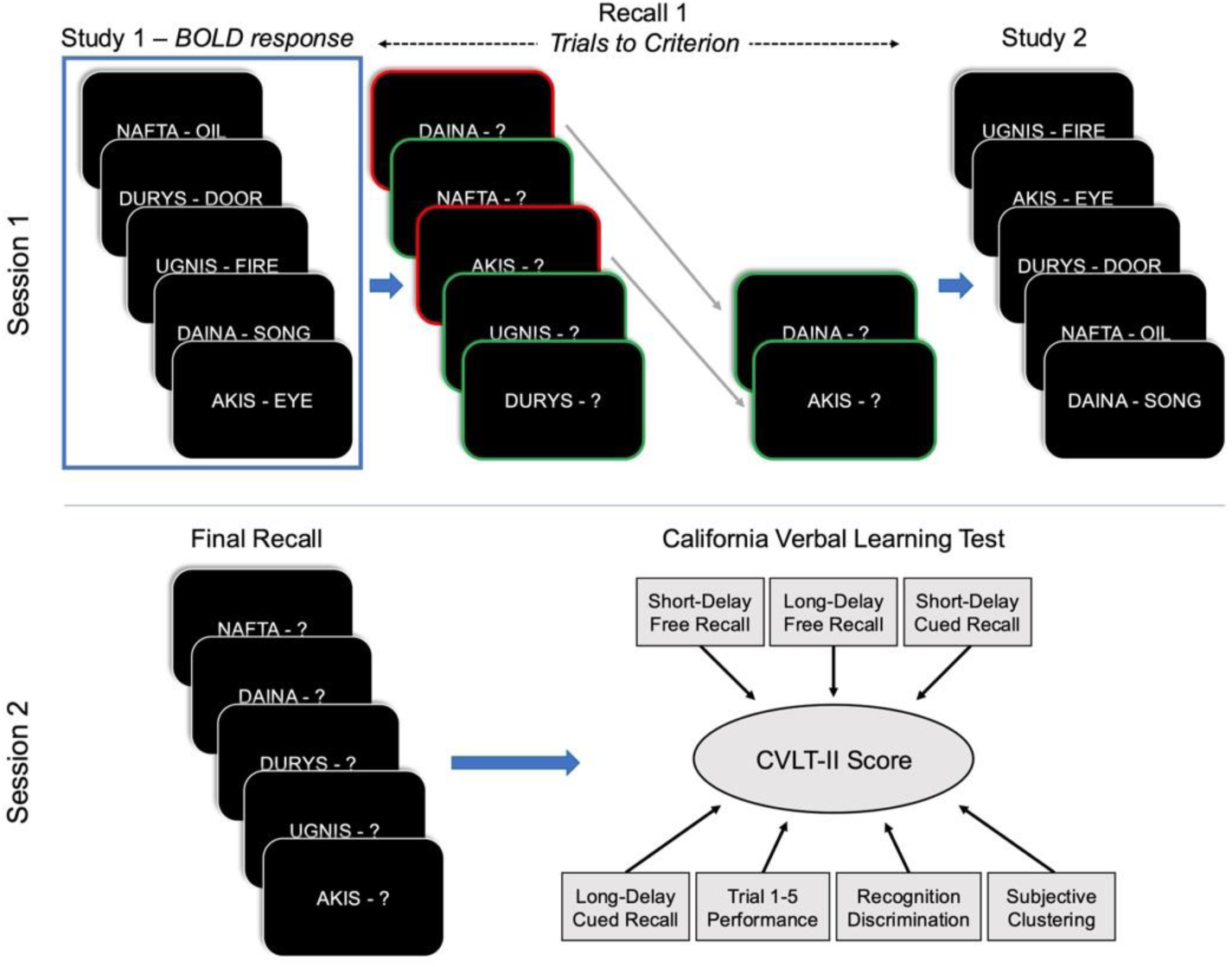
Schematic of the experimental paradigm. In Session 1, participants studied Lithuanian–English word pairs during Study 1, completed iterative cued-recall testing of each word pair until reaching the 100% accuracy criterion during Recall 1, then re-studied the word pairs in Study 2. In Session 2, which occurred 2 days later, participants completed a single iteration of the cued-recall test followed by a cognitive battery that included the CVLT-II. BOLD response was collected during Study 1, Study 2, and 10-minute resting periods preceding Study 1 and Study 2. The presented analyses use BOLD response during Study 1.

In the second phase, Recall 1, participants remained inside the scanner to complete iterative cued-recall tests with feedback until reaching the 100% recall criterion. On each cued-recall trial, participants were shown a Lithuanian cue word (e.g., AKIS) for 3.5s and were instructed to verbally recall the matching English translation (e.g., “eye”). Regardless of response accuracy, the correct English word was displayed for 1.5s following each Lithuanian cue presentation. All 45-word pairs were presented in random order in the first iteration, and correctly-recalled word pairs were removed from the list of pairs to be tested in future iterations. A 1s interstimulus interval separated each item, and a 30s arithmetic problem solving task separated each iteration of this phase. During the arithmetic task, participants solved arithmetic problems aloud to limit the possibility of working memory maintenance of cue-target pairs from the previous iteration. Participants completed this cued-recall test in repeated iterations, separated by arithmetic blocks, with removal of correctly-recalled word pairs, until participants had correctly recalled the English translation of each Lithuanian word cue.

The third phase, Study 2, was identical to Study 1. Both Study 1 and Study 2 were preceded by 10-minute resting-state scans with participants fixated on a crosshair at the center of the screen.

The fourth and final phase, Final Recall, took place 2 days (*range*: 36.5-49.5h, *mean*: 43h) after the previous phases. During this Final Recall phase, participants were presented with Lithuanian cue words for 8s and were instructed to verbally recall the matching English translation. No feedback was provided. All 45 Lithuanian–English word pairs were tested in a single iteration in random order with 1s interstimulus intervals. Following Final Recall, participants completed a cognitive battery including the California Verbal Learning Test (CVLT-II) memory evaluation.

### fMRI Data Acquisition

A standard set of imaging protocols guided acquisition of functional MR images. Foam pillows and a thermoplastic mask fastened to the head coil stabilized participants’ head position. All images were acquired using a Siemens MAGNETOM Tim Trio 3.0T Scanner (Erlangen, Germany) and a Siemens 12 channel Matrix Head Coil. A T1-weighted sagittal Magnetization-Prepared Rapid Gradient-Echo structural image was obtained (time echo [TE] = 3.08 ms, time repetition [TR] partition = 2.4 s, time to inversion [TI]=1000ms, flip angle=8°, 176 slices with 1×1×1mm voxels) [9]. In addition, a T2-weighted turbo spin echo structural image (TE = 84 ms, TR = 6.8 s, 32 slices with 2 × 1 × 4 mm voxels) was obtained in the same anatomical plane as the subsequent BOLD images to improve alignment to an atlas. An auto-align pulse sequence protocol provided in the Siemens imaging software package was used to align the acquired slices from functional scans in parallel to the anterior commissure-posterior commissure plane and centered on the brain. This plane parallels the slices in the Talairach atlas [10], which is used for subsequent data analysis. Functional imaging was performed using a BOLD contrast sensitive gradient echo echo-planar sequence (TE = 27 ms, flip angle = 90°, in-plane resolution = 4 × 4 mm). Whole-brain Echo Planar Imaging volumes of 32 interleaved, 4-mm-thick axial slices were obtained every 2.5 s. The first 4 image acquisitions were discarded to allow net magnetization to reach steady state.

Noise-cancelling headphones were used to help dampen scanner noise for participants. The headset was equipped with a microphone and allowed participants to communicate with research technicians throughout the scanning procedures. An Apple iMac computer (Apple, Cupertino, CA, USA) running PsyScope software [11] and Adobe Flash Professional CS5.5 (Adobe Systems, San Jose, CA, USA) was used to display visual stimuli. An LCD projector (model PG-C20XU, Sharp) was used to project stimuli onto an MRI-compatible rear-projection screen (CinePlex) at the head of the scanner bore. Subjects viewed this screen through a mirror mounted to the top of the head coil.

### fMRI Data Preprocessing

Each subject’s fMRI data was preprocessed with a standardized stream meant to reduce noise and remove artifacts from the data. This protocol included: 1) correction for movement within and across runs using a rigid-body rotation and translation algorithm [12]; 2) mode-1000 intensity normalization, allowing for comparisons across subjects [13]; and 3) temporal realignment of all slices to the temporal midpoint of the first slice using sinc interpolation to account for the slice-time acquisition differences. Functional data were then resampled into 3-mm isotropic voxels and transformed to stereotaxic atlas space [10]. To register individuals’ data to an atlas, subjects’ T1-weighted images were aligned to a custom atlas-transformed [14] target T1-weighted template (711-2B) using a series of affine transforms [15,16].

### Administration of Memory Evaluation

The CVLT-II was administered amongst several other cognitive measures. For additional details regarding other measures, see Nelson et al. [7]. The CVLT-II consisted of two parts. In Part 1, the experimenter read a list of 16 words, and the participant was asked immediately after to recall as many of the words as they could remember in any order. This comprised List A Immediate Free Recall Trial 1 through List A Short Delay Cued Recall [17]. In Part 2, which occurred 20 minutes after Part 1, participants were asked to recall in any order as many of the 16 words as they could remember from the list presented in Part 1. This part included List A Long Delay Free Recall through List A Long Delay Yes/No recognition [17]. Instructions were read directly from the CVLT-II packet. Participants did not complete the Forced-Choice Recognition portion of the CVLT-II.

### Statistical Analyses

#### Analysis of Memory Evaluation

Together, Part 1 and Part 2 of the CVLT-II yielded 16 separate memory measures. A factor analysis of CVLT-II memory measures identified seven measures that loaded onto the first component and accounted for 40.82% of the total variance: “short-delay free recall,” “long-delay free recall,” “short-delay cued recall,” “long-delay cued recall,” “trial 1-5 performance,” “subjective clustering,” and “recognition discrimination.” We z-scored these measures and averaged them for each participant to obtain the CVLT-II score used here. See Nelson et al. [7] for a full list of the CVLT-II memory measures and more details regarding the factor analysis.

#### BOLD Activity Analyses

We subtracted BOLD activity during the Lithuanian word learning task from BOLD activity at rest to obtain a measure of task-induced activity. Our *a priori* replication analysis calculated the BOLD signal change within the exact same region, the right pHPC, in which Wig et al. [6] observed the effect (converted to MNI coordinates: 22, –33, –2). We then calculated the correlation between BOLD signal change and performance on the CVLT-II as measured by the z-scored long-term memory score.

In an exploratory analysis, we defined regions in the hippocampus and parahippocampal cortex that were correlated at p <0.05 (two-tailed test) with CVLT-II score. Regions of interest (ROI) were defined as 10-mm diameter spheres around the peak correlation value in the voxel-wise image. BOLD data from these regions of interest were plotted against CVLT-II scores to examine whether outliers existed in the data and for graphical depiction purposes. Correlation values are presented for the purpose of reporting on the strength of the relationship, not to emphasize that the relationship was statistically significant, which would imply circularity.

## Results

Of primary interest was the replication of the correlation between right posterior hippocampal activity and performance in an out-of-scanner learning task similar to that reported by Wig et al. [6]. We therefore calculated the Pearson correlation coefficient between participant *z*-scores on the CVLT-II long-term memory measure (Figure 2A) and BOLD activity, expressed in terms of percent signal change, in an ROI derived from Wig et al. [6], which consisted of a 10-mm diameter sphere centered around MNI coordinates 22, –33, –2 (Figure 2B). CVLT-II z-scores ranged from –2.21 to 1.82, and percent signal changes in the *a priori* pHPC ROI ranged from –0.47 to 0.59. Within this ROI, there was a significant negative relationship between CVLT-II score and BOLD activity relative to rest (*r* = –0.27, *p* = .012), such that participants with lower activation performed better on the CVLT-II (Figure 2C).

**Figure 2.**
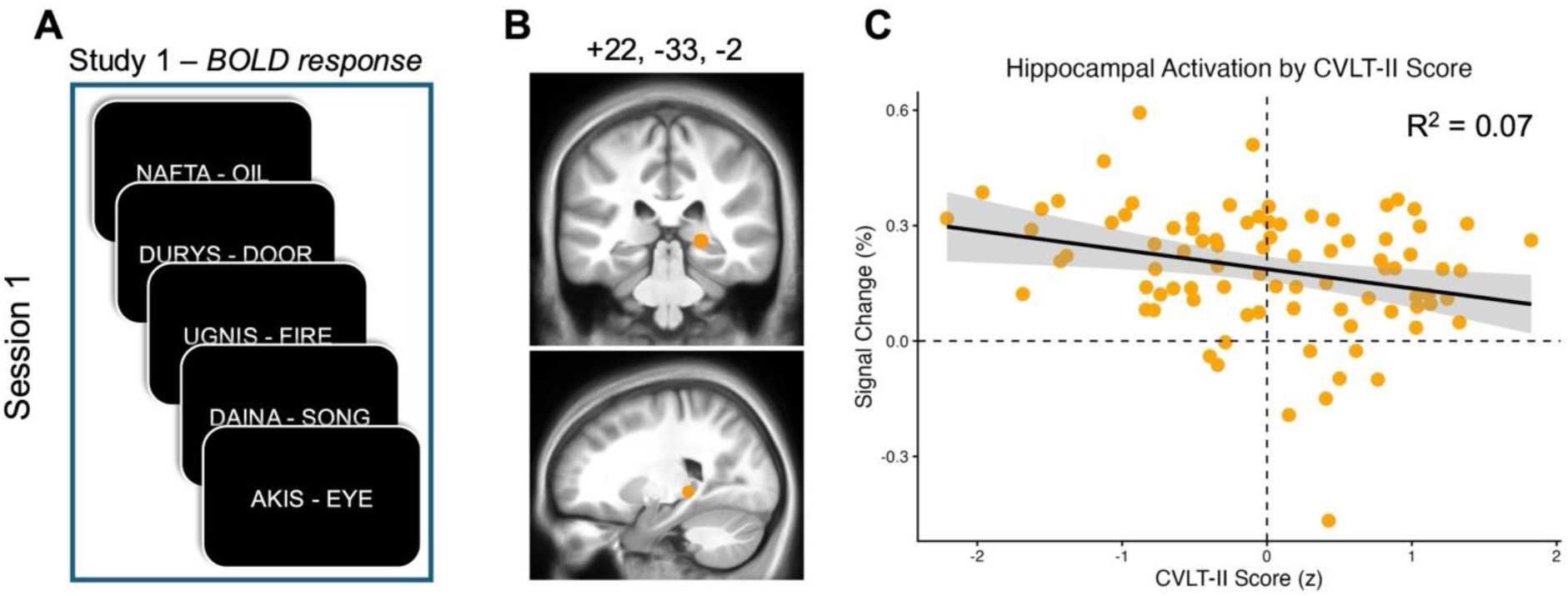
Hippocampal activity correlates with long-term memory performance. (A) Schematic of the task for which we calculated signal change and correlations with CVLT-II performance. (B) Visual representation of the *a priori* brain region of interest in the posterior hippocampus (pHPC), projected onto an MNI152 brain, that was adapted from Wig et al. [6]. MNI coordinates for the region are included above the visual representation. (C) Scatterplot depicts the relationship between CVLT-II score (z-score) and signal change in the right hippocampal tail from rest to Study 1 periods.

These findings represent a near-direct replication of the findings presented in Wig et al. [6], in which hippocampal activity during task performance correlated negatively with a memory component score on a separate memory battery. This was true across two experiments with two different tasks and organizations of resting periods (Figure 3).

**Figure 3.**
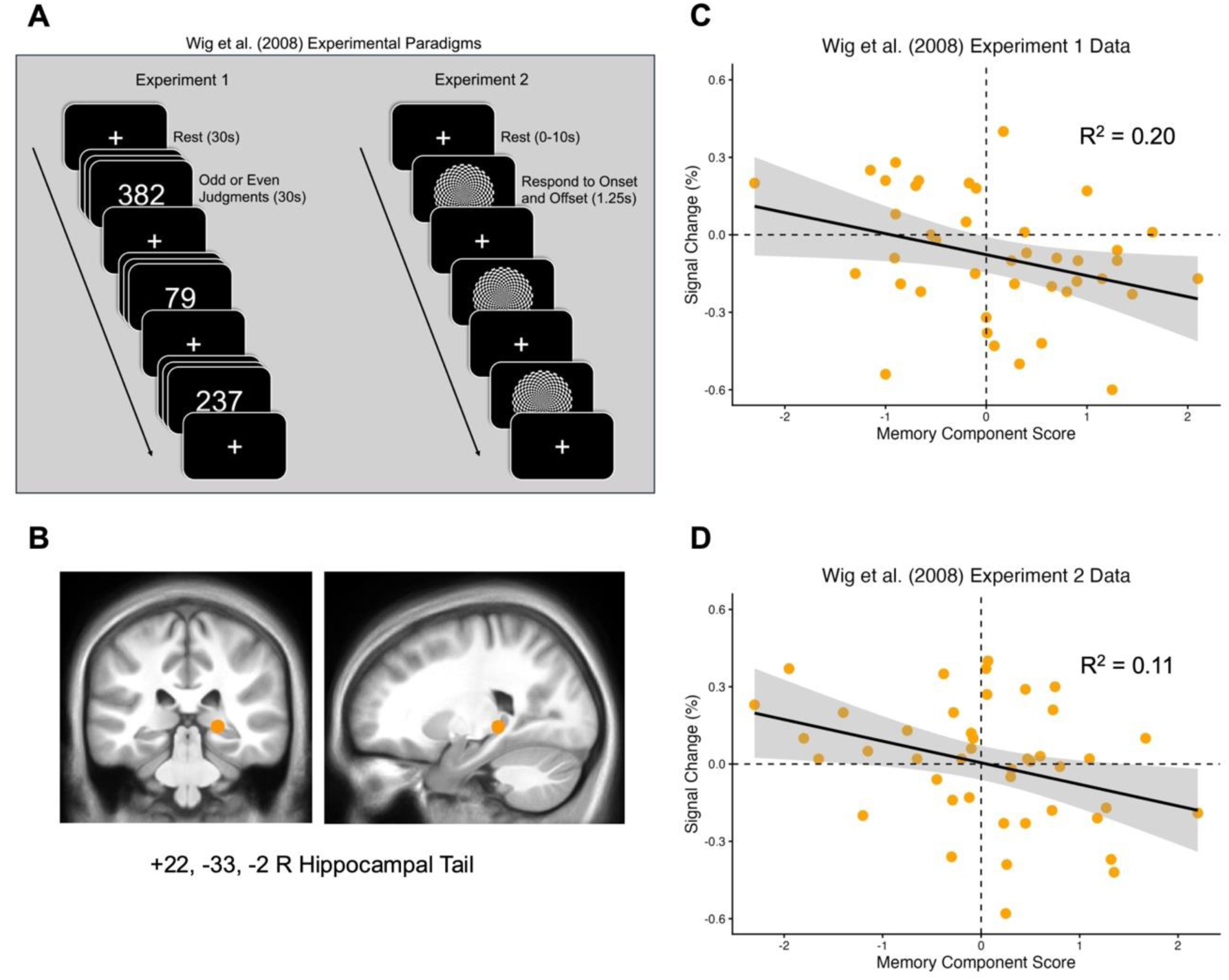
Previously reported data showing a negative relationship between task-induced hippocampal activation relative to rest and performance on an outside memory battery. (A) Schematics of the experimental paradigms used in Exp. 1 and Exp. 2 of Wig et al. [6]. In Exp. 1, participants alternated between 30s resting state periods in which they fixated on central crosshairs and 30s periods in which they made odd or even number judgments. In Experiment 2, participants indicated the onset and offset of a flickering checkerboard with jittered periods of 0-10s of rest between active trials. (B) Depiction of the Wig et al. [6] ROI on an MNI152 brain. (C) Scatterplot of the relationship in Exp. 1 between memory component score and percent signal change in the right hippocampus during the odd or even number judgment task relative to resting periods. (D) Scatterplot of the relationship in Exp. 2 between memory component score and percent signal change in the right hippocampus during the flickering checkerboard task relative to resting periods.

After replicating the right pHPC correlation results from Wig et al. [6], we wanted to better understand the degree to which the observed relationship was specific to this specific hippocampal region. We therefore conducted an exploratory analysis, focusing on the medial temporal lobe (MTL) in which we could identify voxels whose BOLD activity during the Lithuanian-English task was significantly correlated with out-of-scanner behavior. Our analysis identified three regions of the MTL in which task-induced BOLD signal change negatively correlated with CVLT-II performance. These included left perirhinal/entorhinal cortex (*r* = –0.36, *p* < .001), the right parahippocampal gyrus (*r* = –0.37, *p* < .001), and the left pHPC (*r* = –0.39, *p* < .001) (Figure 4). These results suggest that the right pHPC correlation, while reliable, is part of a broader pattern observable across the MTL that links in-scanner activation levels to out-of-scanner performance on a memory task.

**Figure 4.**
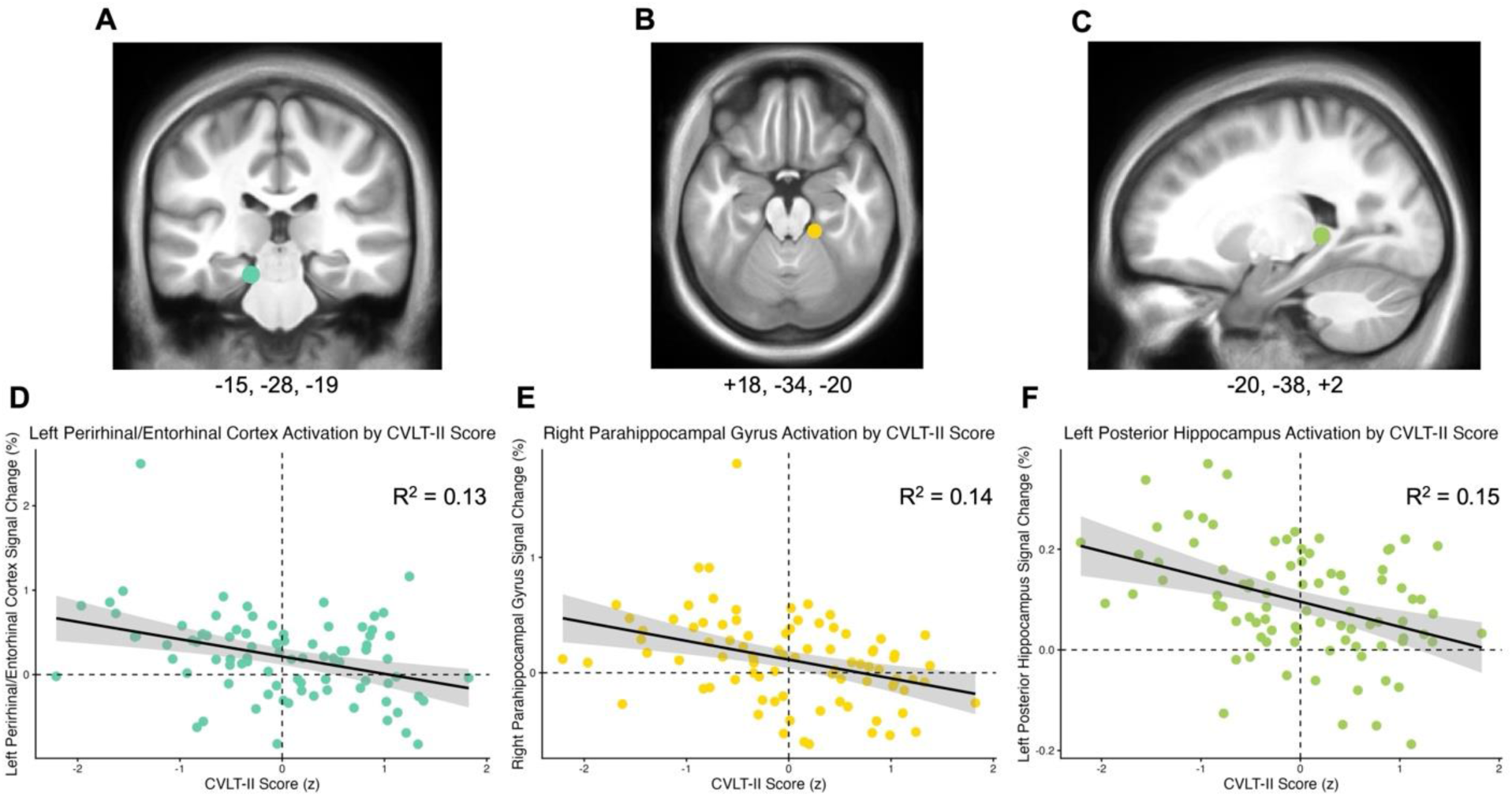
Brain regions with activity that negatively correlates with CVLT-II performance. (A-C) Visual depictions of the brain regions that showed negative correlations with CVLT-II performance. (D-F) Scatterplots showing relationship between BOLD signal change and CVLT-II z-score. Specifically, this exploratory analysis identified correlations with BOLD activity in the left perirhinal/entorhinal cortex (A, D), right parahippocampal gyrus (B, E), and left posterior hippocampus (C, F). MNI coordinates for each region are included under the corresponding visual depiction.

## Discussion

Lower levels of pHPC activation during a verbal paired associates learning task predicted better performance on CVLT-II scores obtained outside of the scanner. These results, obtained in an independent sample of 86 young adults, serve as a clear replication of prior results identifying task-evoked pHPC activity as a predictor of individual differences in verbal memory function. However, when expanding our focus beyond right pHPC, we found that this result is part of a broader pattern that can be observed across regions of the MTL.

A primary purpose of the present investigation was to provide a critical out-of-sample replication to form a basis for future investigations of individual differences in memory ability relating to hippocampal activation levels. That significant effects were observed in the same region, and in the same direction, as the initial findings of Wig et al. [6] suggests that recently raised concerns regarding the need for thousands of subjects in studies of individual differences in brain-behavior correlations [18] may be overstated. Rather, a strong *a priori* prediction, coupled with a task design that allowed for intersubject variability to be clearly observed, allowed for an unambiguous interpretation with fewer than 100 participants. Instead of calling for skepticism, we instead argue that the findings reported here, combined with other recent examples of biologically plausible correlations observed across subjects [19], instead support a more common sense position: the amount of data needed depends greatly on the nature of the question being asked. In this particular case, the results also reinforce the need to further explore the mechanisms that associate pHPC activity and verbal memory, which would include testing for offline consolidation, as proposed by Wig et al. [6]. We also present the first step toward further exploration in the form of an exploratory voxel-wise association analysis. CVLT-II performance was negatively associated with activation during the Lithuanian word learning task in the left perirhinal/entorhinal cortex, right parahippocampal gyrus, and left pHPC. This demonstrates that the negative relationship between task-induced activation and out-of-scanner memory performance is not limited to just the right pHPC region identified in Wig et al. [6] and replicated in our *a priori* analyses. The effect extends to other regions of the MTL as well.

### Hippocampal Markers of Individual Differences in Memory Ability

The present findings complement a large body of previous research demonstrating close relationships between regions of the hippocampus and individual differences in memory performance or capacity. At least three lines of previous research, considered alongside the present findings, support the conclusion that lower task-induced pHPC activation, relative to rest, supports verbal memory ability. First, structure-behavior associations between the hippocampal tail and memory performance highlight pHPC as a region of importance in verbal memory function [20–23]. These structural results converge not only on the *a priori* pHPC ROI initially explored in this report, but also with one of the 3 MTL ROIs subsequently identified in our whole-brain analysis. Second, functional links between pHPC and functional networks involved in cognitive control and familiarity detection [24,25] further emphasize a role for pHPC, specficially, in memory-related cognition and task engagement more generally. Third, the associations between HPC and verbal memory observed in this report and in Wig et al. [6] are consistent with negative correlations observed between in-scanner and out-of-scanner memory performance in earlier work [26]. As an extension of several previous investigations and a near-direct replication of one, our observation that reduced task-induced pHPC activation differentiated individuals of differing memory capabilities establishes this finding as reliable and sufficiently consistent as to support further investigation.

### Hippocampal heterogeneity

Recent reports focusing on functional heterogeneity across regions of the hippocampus provide important insights into how the current results should be interpreted. Across multiple studies using precision MRI approaches in densely sampled individual participants, aHPC and pHPC have been associated with distinct functional networks that support differing forms of memory retrieval [24,25]. The aHPC is strongly functionally connected to the regions of the default mode [27,28] and contextual association [29] networks, and was found to support scene construction processes in an episodic project task [24]. In contrast, the pHPC is functionally connected to the parietal memory network and frontoparietal control network [25] or salience network [24]. Rather than tracking scene construction demands, pHPC appeared to support salience or novelty detection and general task engagement [24], as is consistent with the networks to which it appears most strongly connected. Differences in general pattern of aHPC/pHPC connectivity with broader functional networks have also been viewed in larger group level analyses, offering further support to the recent precision mapping results [30–33]; see also [4].

With respect to the current results, pHPC, but not aHPC, regions emerged across both targeted ROI and follow-up exploratory analyses. If, as prior evidence suggests, pHPC activity tracks task engagement or salience [24] and supports the processing of fine-grained details in the context of episodic memory [4,34], differences in task-induced perturbations across participants could be viewed as reflecting general mnemonic information processing efficiency, rather than reflecting something like post-encoding rehearsal that would involve the recollective or scene construction processes associated with aHPC and DMN regions. This account differs from the consolidation emphasis originally posited by Wig et al. [6] and is a promising target for more specific analyses given the reliability of the correlation being observed.

### Voxelwise analysis of in-scanner BOLD and out-of-scanner task performance correlations

In addition to replicating the Wig et al. [6] results, which was a key motivator of the current experiment, the data also allowed for an assessment of the specificity of the effect in the right pHPC. Voxelwise results highlighted that the effect observed in right pHPC is part of a broader pattern of MTL regions that exhibited the same overall pattern: reduced task-induced activity correlating with better offline performance. These regions included the left perirhinal/entorhinal cortex, right parahippocampal gyrus, and, as noted previously, a homotopic left pHPC region to accompany the right pHPC carried forward from Wig et al. [6]. Parahippocampal and perirhinal cortex receive inputs that emphasize spatial/contextual and item-specific information, respectively [35,36], and in turn are connected to the hippocampus via the entorhinal cortex [37]. Here, as with the functional connectivity results discussed above, past findings support an account by which the correlations observed in these data reflect broad efficiency differences general mnemonic information processing (and further suggest that the underlying mechanism need not be hippocampus-specific, which can be better determined in follow-up work).

### Limitations

Although the present study represents a necessary replication and extension of previously reported findings, there are important limitations to consider. First, the nature of our targeted replication led us to limit *a priori* analyses of brain-behavior correlations to a single region within pHPC. While benefitting from strong theoretical motivation, it was necessarily limited to a small portion of a single structure. Our follow-up voxel-wise analysis allowed for a broader exploration of neural associates of individual differences in memory behavior that helped to determine the degree to which the present findings are exclusive to pHPC, but as a trade-off, was more limited in its statistical power. In principle, it also became and more susceptible to the pitfalls of brain wise association studies that often lead to inflated effect sizes [18]. Although the consistency of the direction and approximate magnitude of the voxelwise effects offers some reassurance, the precise correlation magnitudes should be interpreted with this possibility in mind.

A related limitation is that the *a priori* pHPC region, combined with the voxel size of the acquired data, did not allow us to localize effects to specific hippocampal subfields, which have been shown to exhibit different effects in memory tasks [38,39]. Similarly, we could not confidently attribute the rhinal cortex pattern specifically to either entorhinal cortex or perirhinal cortex, despite the functional differences known to exist between these structures [40]. Future experiments, perhaps conducted at 7T and with sub-millimeter voxel sizes, would offer a more specific view of the basis for the observed individual differences within the hippocampus specifically and the MTL more broadly.

We calculated task-induced activation levels against a resting state baseline. Although there is evidence that it constitutes an interpretable baseline that aligns with neural baseline measures and metabolic baseline as established by PET [27,41], others have argued that its unconstrained nature poses difficulties because the hippocampus is always actively encoding and retrieving information [39,42,43]. In contrast, foil task baselines constrain cognitive activity more overtly and, if multiple baseline tasks are employed, would provide an opportunity to isolate task components that are differentially correlated with pHPC activity across. Indeed, one previous study that used an active foil task as baseline also localized individual differences in CVLT-II performance to the hippocampus [26], providing convergent support for our results.

### Conclusions

In sum, our findings converge with an earlier body of work showing a close connection between the hippocampus and individual differences in memory-related cognition. By identifying a replicable negative correlation between task-induced hippocampal activity and scores on a general memory evaluation and extending this finding to other closely related regions of the MTL, the present findings indicate the pHPC-verbal memory relationship is both consistent and reliable. By externally replicating and then extending Wig et al. [6], the present findings now provide a motivation for future research into the nature of why task-induced pHPC changes reflect individual differences, particularly in the counter-intuitive direction that was observed, and how this might be used to better understand expression and risk of memory impairment.

